# Giant virus transcription and translation occur at well-defined locations within amoeba host cells

**DOI:** 10.1101/2025.03.24.645094

**Authors:** Lotte Mayer, Georgi Nikolov, Martin Kunert, Matthias Horn, Anouk Willemsen

## Abstract

Many giant viruses replicate in the cytoplasm in so-called viral factories. How exactly these viral factories are established and where the different steps of the replication cycle occur remains largely obscure. We have developed a single-molecule messenger RNA fluorescence in situ hybridisation (smFISH) protocol for giant viruses in an *Acanthamoeba* host. Combined with other labelling techniques (FUNCAT, DiD, rRNA FISH, DAPI), we show the Mimivirus transcription and translation sites during an infection cycle in the amoeba host cell. While viral mRNA localisation changes depending on the infection stage, transcription occurs at well-defined spots within the viral factory. The original viral cores released within the cytoplasm most likely define these spots. When transported outside of the viral factory, the translation of viral mRNA takes place in a well-defined ring surrounding it. With this study, we obtained novel insights into giant virus replication, of which the methods are widely applicable to other viruses for the visualisation and quantification of RNA molecules.

## Introduction

Most DNA viruses carry out their replication and transcription either entirely or partially within host nuclei. While the nuclear environment can considerably enhance the efficiency of replication and transcription (Whittaker et al., 2000), it also presents several hurdles, as viral molecules need to be transported to and into the nucleus. Large DNA viruses are particularly affected by hurdles associated with genome trafficking, as the mobility of DNA molecules in the cytoplasm and the nucleus is size-dependent (Lukacs et al., 2000), with the main barrier for larger molecules being the cytoskeleton of the host cells (Dauty & Verkman, 2005).

Large DNA viruses belonging to the phylum *Nucleocytoviricota* were previously referred to as nucleocytoplasmic large DNA viruses (NCLDVs). As the name suggests, these viruses can replicate in the host cell nucleus and/or cytoplasm (Koonin et al., 2019). Before the discovery of so-called giant viruses in 2003 (La Scola et al., 2003; Raoult et al., 2004), the only other large DNA viruses that were known to have an entire cytoplasmic infection cycle were the poxviruses (Minnigan & Moyer, 1985; Schramm & Locker, 2005). The cytoplasmic life cycle of poxviruses can be roughly divided into virion entry, early transcription, DNA replication, followed by virus assembly and release. It has been recently shown that the giant Mimivirus displays extensive physiologic similarities with poxviruses (Mutsafi et al., 2010, 2013). The Mimivirus genome is not delivered to the host nucleus and never crosses the nuclear membrane (Mutsafi et al., 2010). A typical Mimivirus infection starts with uptake by the amoeba host cell through phagocytosis. Within the phagosome, the capsid of Mimivirus is opened through a so-called stargate, which allows for the fusion of the viral and the phagosome membrane, forming a large tube (Zauberman et al., 2008). Through this membrane tube, the genome core is released (possibly within a vesicle) into the host cytoplasm. Shortly after this delivery, transcription is initiated within the Mimivirus core, followed by a burst of DNA replication within a cytoplasmic viral factory surrounding the core (Mutsafi et al., 2010). DNA packaging into preformed Mimivirus procapsids proceeds through a portal that is transiently formed at a site distal to the stargate (Zauberman et al., 2008). Therefore, despite their distant phylogenetic positions (Aylward et al., 2021), poxviruses and Mimivirus appear to have evolved similar mechanisms to cope effectively with the exit and entry of large genomes.

Due to the large particle and viral factory sizes of most giant viruses (*e.g.* mimiviruses and marseilleviruses), we can observe infections in single host cells using general nucleic acid staining (*e.g*. DAPI) combined with fluorescence microscopy. However, this method is not virus-specific and is limited to those giant viruses that form viral factories (*e.g.* medusaviruses and pandoraviruses do not form viral factories). In this study, we have developed a single-molecule messenger RNA fluorescence in situ hybridisation (smFISH) protocol that allows us to study active giant virus infections independent of their replication strategy. Combined with other techniques to label DNA, proteins, and membranes, rRNA FISH of the host, and digital PCR we learn more about the viral transcription and translation sites and demonstrate that the methods presented here have ample applications for studying giant virus infections within amoeba host cells.

## Results and Discussion

### Setting up smFISH for giant viruses

The single-molecule messenger RNA fluorescence in situ hybridisation (smFISH) technique identifies a single mRNA based on the binding of multiple small probes targeted to different locations on the mRNA of interest (Aagaard et al., 2017; Raj et al., 2008). To set up smFISH for giant viruses within *Acanhtamoeba* host cells (Fig. 1), we designed and synthesised a set of 35 individually fluorophore-labelled probes that target the major capsid protein (*mcp*) gene of Mimivirus (Materials and Methods, Table S1). The *mcp* gene is expressed at the late stage of a typical Mimivirus infection, between 6 and 12 hours post-infection (h.p.i) (Legendre et al., 2010). In contrast, no *mcp* expression was detected during the early (< 3 h.p.i) and intermediate (> 3 h.p.i and < 6 h.p.i) stages of infection (Legendre et al., 2010). Different conditions for fixation and permeabilisation of the amoeba cells, hybridisation of the probes, washing temperature and concentration of the probes were tested (Materials and Methods) to obtain the best signal with the optimal conditions in Table 1.

**Figure 1.**
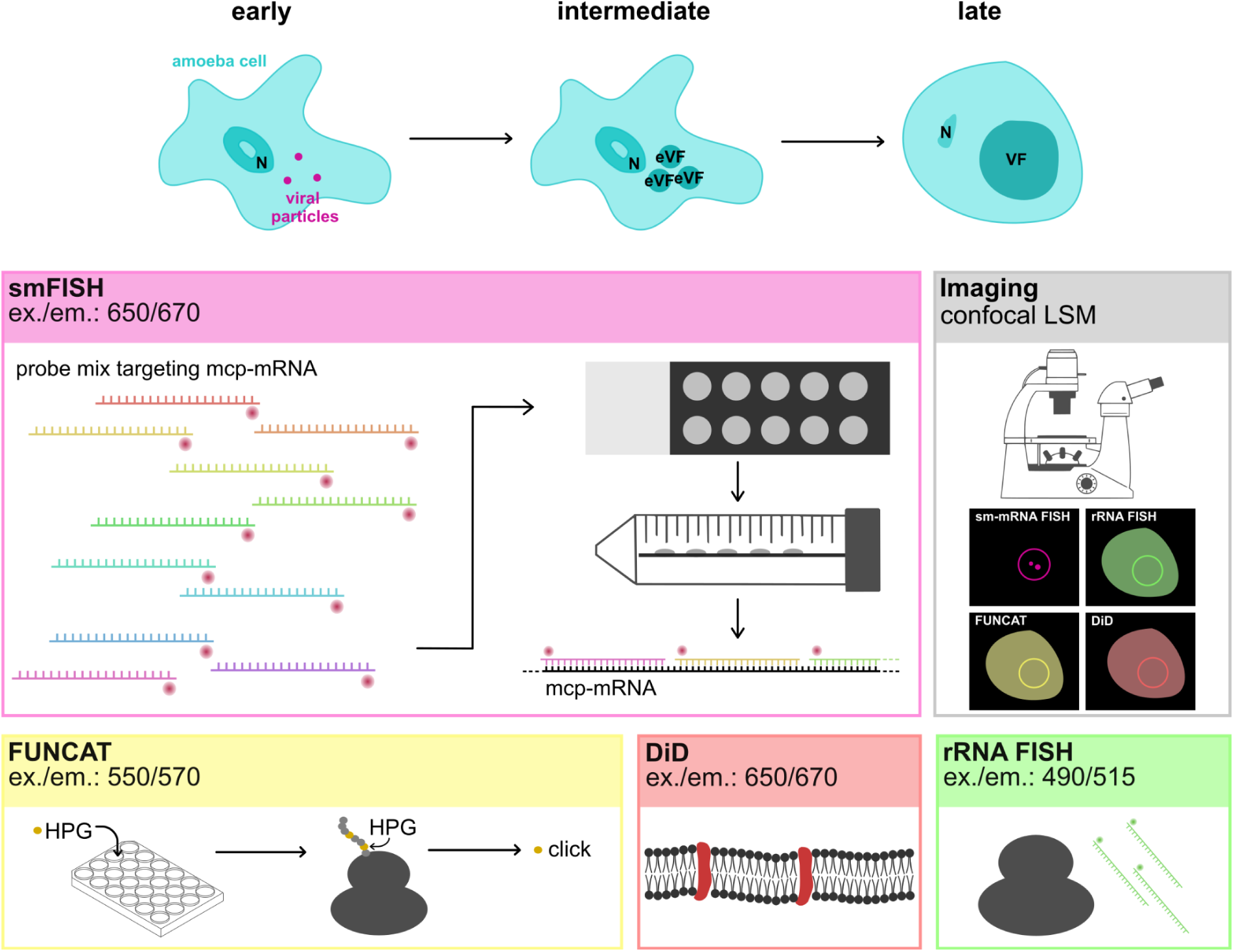
An overview of the methods used in this study. The top of the image shows different infection stages of Mimivirus within an amoeba cell, where the cell gets infected (early: < 3 h.p.i), where early viral factories (eVFs) are formed within the cell (intermediate: > 3 h.p.i and < 6 h.p.i), and where the eVFs have fused into one mature VF (late: > 6 h.p.i). To study these infection stages, we use single-molecule messenger RNA fluorescence in situ hybridisation (smFISH), fluorescent noncanonical amino acid tagging (FUNCAT), DiD (1,1′-dioctadecyl-3,3,3′,3′-tetramethylindodicarbocyanine, 4-chlorobenzenesulfonate salt) labelling, and ribosomal RNA (rRNA) FISH. The fluorescence of these labels is subsequently visualised using confocal laser scanning microscopy (CLSM).

**Table 1.** Optimal conditions for smFISH targeting the *mcp* gene of Mimivirus.

|  |  |  |
| --- | --- | --- |
| Sample preparation | Fixation | 4% formaldehyde, 10 min, RT |
|  | Permeabilisation | 70% Ethanol, 1 hour, 4°C |
| Hybridisation | Ingredients (final concentration) | 4.375 ng/μL probe mix |
|  |  | 20% formamide |
|  |  | 1 mg/mL *E. coli* tRNA |
|  |  | 2X SSC (saline-sodium citrate) |
|  |  | 0.2 mg/mL BSA (bovine serum albumin) |
|  |  | 2 mM Ribonucleoside-vanadyl complex |
|  |  | 100 mg/mL Dextran sulfate |
|  |  | Nuclease free water |
|  | Temperature | 37°C |
|  | Time | Overnight |
| Post-hybridisation | Initial wash | 20% formamide, 2X SSC, 15 min 39°C |
|  | Final wash | Ice-cold Milli-Q water |

Different percentages of formamide, ranging from 0% to 50%, were tested in the hybridisation buffer. These formamide series show that while 10% formamide already gives a crisp signal (Fig. S1), the optimal formamide concentration for this probe mix and target sequence is 20%, as this concentration leads to less variability in fluorescent signals between experiments. As a negative control, the cells were treated with RNAse A before fixation. No signal was observed when performing the same protocol on RNAse-treated cells (Fig. S2).

### Mimivirus mcp mRNA localisation changes during the infection cycle

By combining electron tomography and fluorescence microscopy, it has been previously shown that each Mimivirus core delivered to the host cytoplasm forms a replication factory and that multiple early viral factories (eVFs) within one host cell can fuse to become one mature viral factory (VF) at late infection stages (Mutsafi 2010). By using bromodeoxyuridine (BrdU) labelling, it was shown that Mimivirus DNA replication occurs in the host cytoplasm after the exit of the Mimivirus genomes from the viral cores (Mutsafi et al., 2010). The same study suggests that early Mimivirus transcription is initiated in the cores during DNA release, and bromouridine (BrU) labelling shows that the newly synthesised mRNAs accumulate at discrete cytoplasmic sites that are adjacent to the DNA replication sites (Mutsafi et al., 2010). To better understand the intracellular localisation of Mimivirus *mcp* mRNA during infection, we use the newly developed smFISH method.

*Acanthamoeba terricola* Neff host cells were infected with Mimivirus, and cells were harvested at different time points during the infection cycle. At the intermediate infection stage of 4 h.p.i, we observe multiple eVFs within each host cell that later fuse (> 6 h.p.i) to become mature VFs (Fig. 2, Fig. S3-S4). Consistent with transcriptomic data (Legendre et al., 2010), we do not observe Mimivirus *mcp* mRNA at 4 h.p.i (Fig. 2B, Fig. 3A, Fig. S3). The earliest time point where we observe *mcp* mRNA is at 6 h.p.i (Fig. 2E), whereas the latest time point is between 18-24 h.p.i (Fig. 2H), right before host cell lysis. Interestingly, at these late infection stages, the intracellular location of mRNA varies at different time points. At 6 h.p.i, when the mature VF has been recently established, the *mcp* mRNA is located mainly in a ring surrounding the viral factory (Fig. 2E). However, at 20 h.p.i, the *mcp* mRNA is also situated at discrete sites within the viral factory (Fig. 2H, Fig. 3A, Fig. 4B, Fig. S2-3), indicating that late Mimivirus *mcp* transcription takes place here. These discrete sites appear as holes within the strong DAPI signal of the VF. Within these holes, the *mcp* mRNA signal is present (Fig. S5), where phase separation (Rigou et al., 2024) is likely responsible for maintaining these sites.

**Figure 2.**
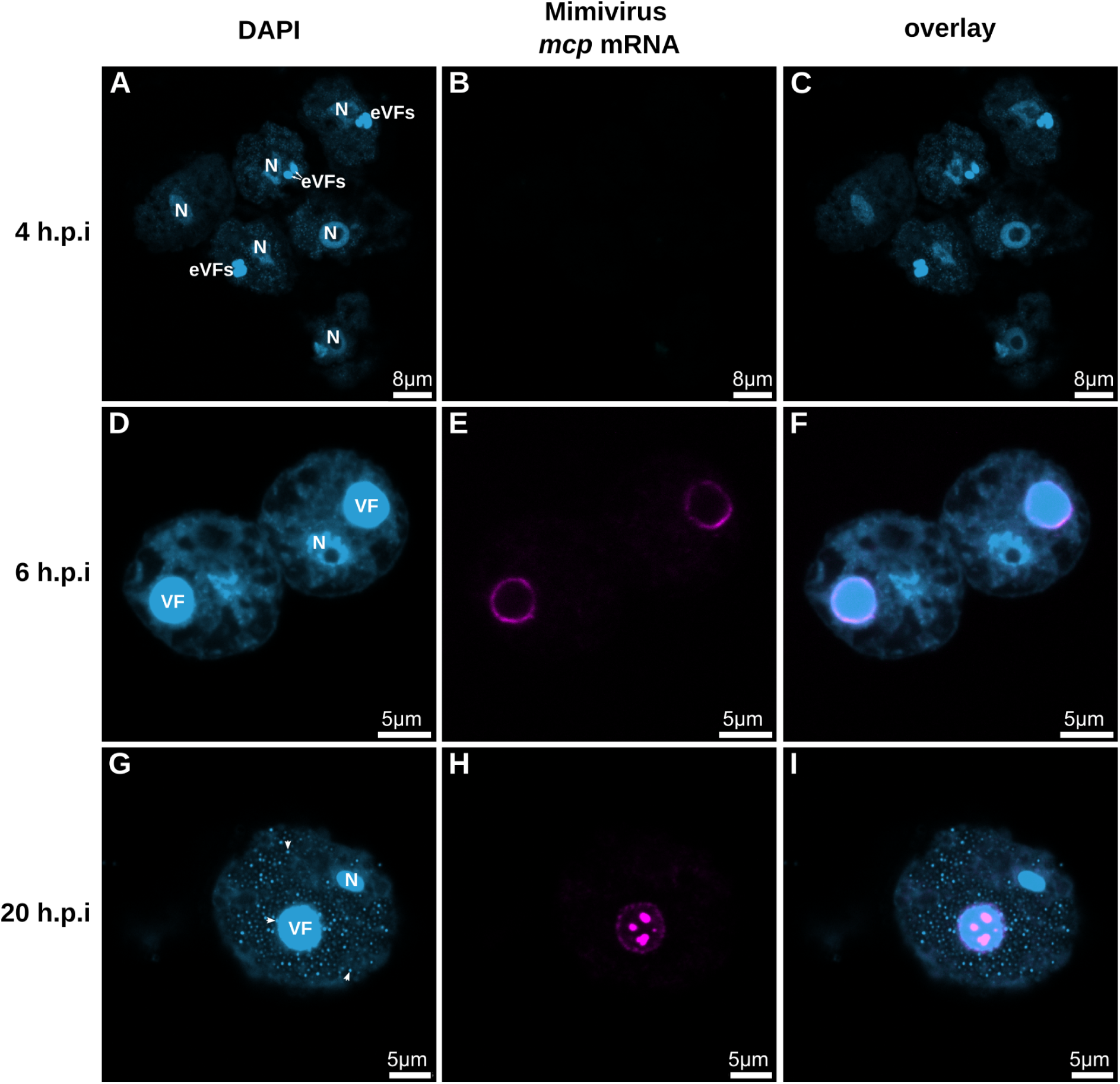
Mimivirus mcp mRNA localises at different sites during the course of infection. (**A**) DAPI staining (light blue) of six amoeba cells, three infected, at 4 h.p.i. A brighter DAPI signal can be observed in the nucleus (N) and the early viral factories (eVFs) of the cells. (**B**) No Mimivirus *mcp* mRNA signal (magenta) is visible at 4 h.p.i. (**C**, **F**, **I**) Overlay of DAPI staining (light blue) and Mimivirus *mcp* mRNA signal (magenta). (**D**) DAPI staining (light blue) of two infected amoeba cells at 6 h.p.i. (**E**) Mimivirus *mcp* mRNA signal (magenta) is localised around the viral factories. (**G**) DAPI staining (light blue) of one infected amoeba cell at 20 h.p.i. A brighter DAPI signal can be observed in the nucleus (N), the viral factory (VF), and the viral particles (white arrows indicate 3 examples) within the cell. The viral particles can be observed budding out of the viral factory and within the rest of the host cell. (**H**) Mimivirus *mcp* mRNA signal (magenta) of one infected amoeba cell at 20 h.p.i. is localised surrounding and at specific spots within the viral factory. The scale bars are drawn on top of the original ones for visibility.

**Figure 3.**
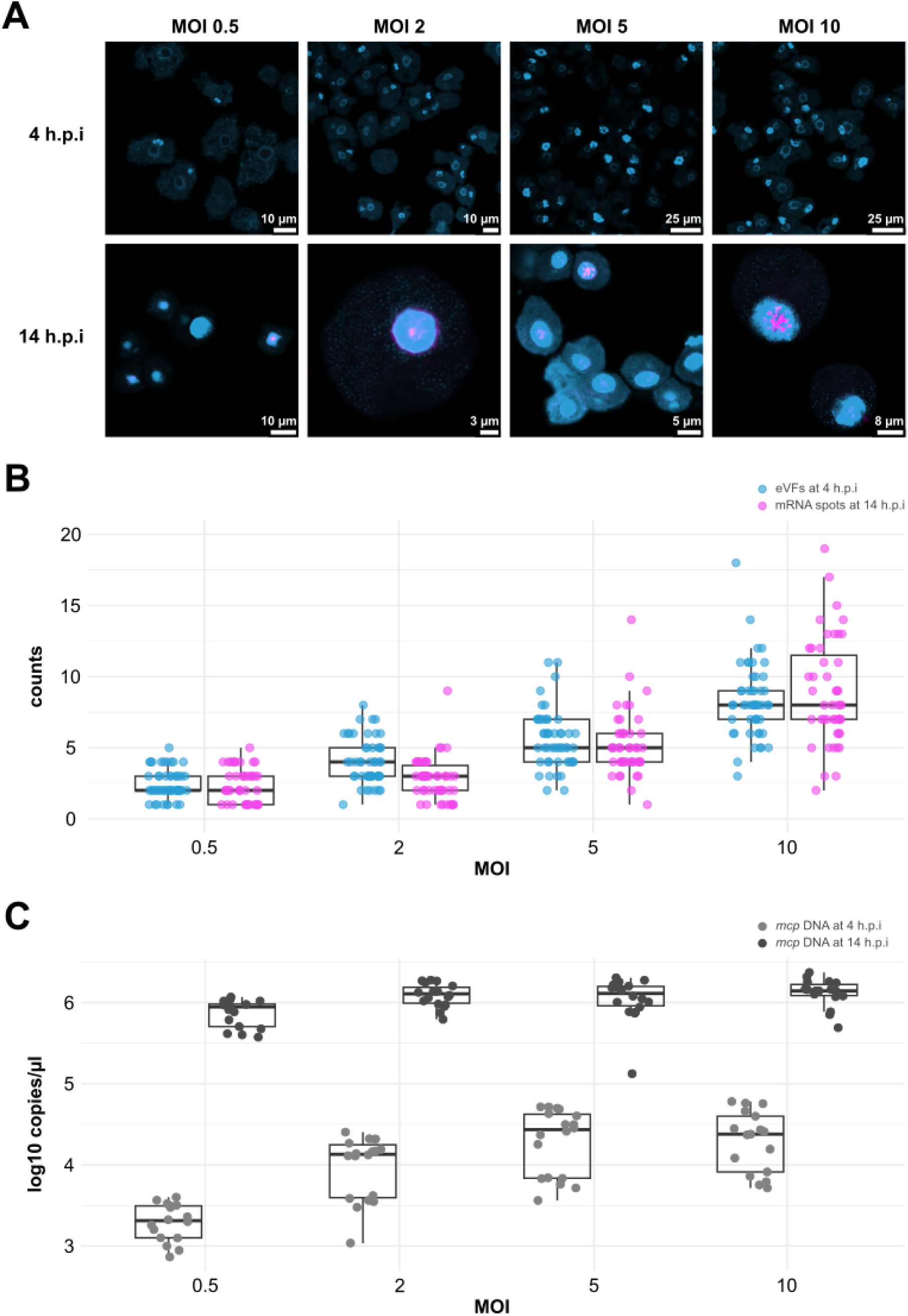
The original viral cores possibly serve separate transcription sites within the fused mature viral factory. (**A**) DAPI staining (light blue) and Mimivirus *mcp* mRNA signal (magenta) of amoeba cells infected with different MOIs at 4 and 14 h.p.i. The Mimivirus *mcp* mRNA signal can not be observed at 4 h.p.i as expression levels are undetectable at this time point (**B**) The number of early viral factories counted at 4 h.p.i and the number of distinct mRNA sites counted at 14 h.p.i after infection with different MOIs. Please note that only infected cells were considered for counting eVFs and mRNA spots. Therefore, the MOI 0.5 and MOI 2 appear inflated compared to the actual MOI used. (**C**) The viral copy number was measured by digital PCR at 4 h.p.i and 14 h.p.i.

**Figure 4.**
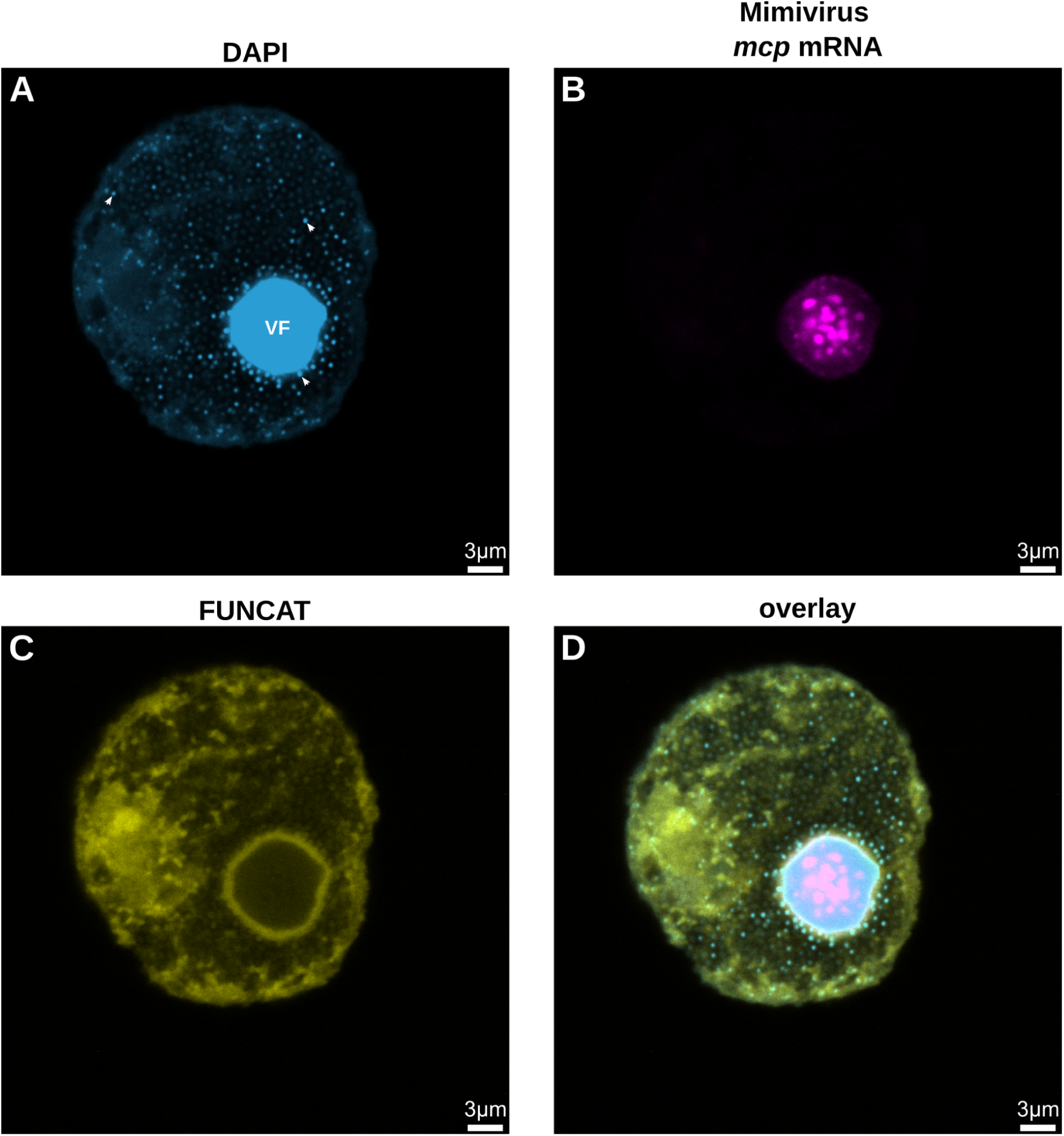
Translation localisation coincides with mRNA localisation surrounding the viral factory at late infection stages. (**A**) DAPI staining (light blue) of one infected amoeba cell at 21 h.p.i. A brighter DAPI signal can be observed in the viral factory (VF) of the cell and for viral particles (white arrows indicate three examples) that can be seen budding out of the viral factory and filling the rest of the cell. (**B**) Mimivirus *mcp* mRNA signal (magenta) of the same cell as in panel A. The mRNA is localised surrounding the viral factories and at specific spots within the viral factories. (**C**) FUNCAT signal (yellow) of the same cell as in panel A. The strong FUNCAT signal suggests that the exported Mimivirus mRNA is translated here. (**D**) Overlay of DAPI staining (light blue), Mimivirus *mcp* mRNA signal (magenta) and FUNCAT (yellow) of the same cell as in panel A. For visibility, the scale bars are drawn on top of the original ones.

While the *mcp* mRNA in this study is detectable at the sites of late transcription, previous research has already shown that early transcription starts in the separate viral cores before fusion (Mutsafi et al., 2010). Therefore, our results suggest that Mimivirus *mcp* mRNA is transported outside the VF at the start of the late infection stage (after fusion of the eVFs), and that upon progression of the late infection stage, more *mcp* mRNA is produced within the centre of the VF. Despite the fusion of the eVFs into one VF, late transcription likely occurs at the same discrete sites where early mRNAs are produced: the remnant sites of the original viral cores.

### The original viral cores in a mature VF may define the location of separate transcription sites

To investigate whether the discrete mRNA spots that we observe at a late infection stage in the mature VF (Fig. 2H) are produced at the remnant sites of the original Mimivirus cores; we followed the course of a Mimivirus infection with an increasing multiplicity of infection (MOI: the number of viral particles per cell).

Consistent with Mutsafi et al., 2010, we observe that with an increasing MOI, a higher number of eVFs are formed within each cell (Fig. 3A-B at 4 h.p.i; *Spearman’s rho* = 0.783, *p-value* < 2.2e-16, *S* = 293682, see also Fig. S3). This increase is significant at each MOI step (Fig. 3A-B at 4 h.p.i and Table S2). Since mRNA of the *mcp* gene of Mimivirus is not detectable at this time point, we counted distinct *mcp* mRNA spots at a later stage of the infection; 14 h.p.i. Also, here, we observe that there is a positive correlation between the MOI and the number of distinct viral mRNA spots present in the fused VFs (Fig. 3B at 14 h.p.i; *Spearman’s rho* = 0.749, *p-value* < 2.2e-16, *S* = 260134). The increase in the counted number of mRNA spots is significant at most steps (Fig. 3AB and Table S2) except for the first (from MOI 0.5 to MOI 2). When comparing the counted eVFs and the distinct mRNA spots, there is no significant difference at most steps, except for MOI 2 (Fig. 3B and Table S2). These results suggest that the original viral cores remain separate transcription sites, even after fusion into one VF. After the fusion of the eVFs, *mcp* mRNA is directly transported outside the VF (Fig. 2E), likely for subsequent translation. Whether this mRNA is produced within the eVFs (between 4 and 6 h.p.i) or in the mature VF right after fusion remains to be investigated.

### More eVFs do not necessarily lead to more viral copy numbers produced by the VF

Together with the increase in MOI, the number of eVFs and the number of mRNA sites, we also observe a positive correlation (Fig. 3C at 4 h.p.i; *Spearman’s rho* = 0.688, *p-value* = 6.428e-11, *S* = 17070) and significant increases in *mcp* DNA copy number at 4 h.p.i up to MOI 5 (Fig. 3C and Table S3), as measured by digital PCR as a proxy for viral copy number. Despite more eVFs being established together with an increase in *mcp* mRNA spots, there is no significant difference in *mcp* DNA copy number between MOI 5 and MOI 10 at 4 h.p.i. This suggests that the maximum space and/or resources for DNA replication are occupied after an MOI of 5 at this intermediate stage of infection. When directly comparing the intermediate and late infection stages, there is a significant increase in DNA copy number at all MOIs (Fig. 3C and Table S3 between 4 and 14 h.p.i). At the late infection stage, the saturation point is reached at MOI 2, as there are no significant differences between MOI 2, MOI 5 and MOI 10 at 14 h.p.i (Fig. 3C and Table S3). Therefore, the differences in MOI mainly affect the DNA copy number at the intermediate stage of infection before the fusion of the eVFs. This observation is backed up by a reported burst of DNA replication during the intermediate infection stage, after the release of the viral genomes into the cytoplasm (Mutsafi et al., 2010). Once fused into mature VFs, the final output in viral copy number is only affected when the MOI is low, most likely because not all cells in the amoeba population are infected. With a high MOI, when all cells are infected (MOI > 1), it does not appear to matter how many viral cores are delivered to the host cytoplasm; the output in viral copy number remains the same. These results suggest a cap on DNA replication within the mature VF.

### Translation of Mimivirus mRNA takes place in a ring surrounding the VF

Our results demonstrate that exported mRNA remains near the VF (Fig. 2E), suggesting that viral translation might occur here. To investigate this, we have combined smFISH with bioorthogonal non-canonical amino acid tagging (BONCAT). This method allows for the identification of newly synthesised proteins (Dieterich et al., 2006; Hatzenpichler & Orphan, 2016; Pasulka et al., 2018) and, when combined with fluorescent tags for visualisation with fluorescence microscopy, is also referred to as fluorescent noncanonical amino acid tagging (FUNCAT) (Dieterich et al., 2010; Iwasaki & Ingolia, 2017).

Indeed, the mRNA ring outside the VF, colocalises with a ring of FUNCAT signal, suggesting active translation of the viral mRNA at this site (Fig. 4). Recently, we described that giant viruses and their known hosts frequently mismatch in codon usage (Willemsen et al., 2024). How giant viruses overcome this mismatch and still ensure efficient viral translation is not clear. For Mimivirus, it seems that the tRNA pool is not significantly altered during the course of infection in their best-known amoeba host (Zhang et al., 2024). This is despite Mimivirus encoding for six tRNA genes. Therefore, we combined smFISH and FUNCAT, with FISH targeting amoebal ribosomal RNA (rRNA), and we show that the rRNA also colocalises in a ring surrounding the VF (Zhang et al., 2024; see also Fig. S4), suggesting that Mimivirus manipulates the translation system to overcome the codon usage mismatch.

When labelled with a lipophilic carbocyanine DiD dye, which enhances fluorescence when incorporated into membranes or bound to lipophilic biomolecules such as proteins, we confirmed the presence of these at the same ring surrounding the VF (Fig. S4). This signal likely indicates the host membranes known to be recruited to the VF (Mutsafi et al., 2013). These host membranes have the characteristic appearance of rough endoplasmic reticulum (ER). They are studded with ribosomes (Mutsafi et al., 2013), supporting the hypothesis that the translation of Mimivirus mRNA occurs at this ring surrounding the VF.

## Conclusion

With the work presented here, we gained novel insights into giant virus transcription, translation and DNA replication. While smFISH has already been used to study active viral infections of a large DNA virus in complex samples (Vincent et al., 2021), it has not yet been used for visualising the localisation of giant virus mRNA. The results in this paper strongly suggest that Mimivirus transcription occurs at the remnant sites of the original viral cores within the fused VF. Recently, it has been shown that the Mimivirus VF is formed by a multilayered phase separation driven by at least two scaffold proteins (Rigou et al., 2024). The presence of proteins associated with transcription in the so-called Inner Layer (a VF compartment analogous to the internal content of the viral core) supports our findings. Moreover, phase separation is likely the driver of maintaining these remnant sites, even after the eVFs are fused into one VF.

Once the Mimivirus cores are delivered within the host cell, early viral transcription starts within these cores. When the viral genome is released from the cores, DNA replication occurs adjacent to these cores/transcription sites (Mutsafi et al., 2010), which leads to the formation of the eVFs. Our results demonstrate that it does not matter how eVFs are present within a single cell; the final output in viral genome copy number remains the same. Whether a mature VF can only produce a maximum number of viral genome copies due to space or resource limitations or some sort of regulation remains to be investigated.

We also show that Mimivirus mRNA is transported outside the VF for translation, which occurs at the border of the VF once it is established. How exactly the mRNA is transported remains to be elucidated. Interestingly, a second Mimivirus VF compartment, the Outer Layer, acts as a selective barrier and recruits VF proteins (Rigou et al., 2024). In a mature VF, the Outer Layer is also localised in a ring around the VF (just like the exported mRNA) and concentrates proteins associated with post-transcriptional regulation. Therefore, we hypothesise that once the mRNA passes the Inner and Outer Layers, it directly encounters the ribosomes on the ER-like host membranes for translation.

Whether the localisation of mRNA of other infection stages is the same and whether the localisation of transcription and translation is shared among all *Nucleocytoviricota* remains to be illuminated. Exciting is the difference between cytoplasmic and nuclear replication viruses. Even though the nuclear viruses transform the host nucleus into a VF, we expect the dynamics to be divergent. Moreover, some *Nucleocytoviricota* encode parts of the translation machinery; therefore, each virus’s genomic information is also likely to have a meaningful impact.

## Materials and Methods

### Probe design

Custom Stellaris® FISH Probes were designed against the *Acanthamoeba polyphaga mimivirus* (GenBank: HQ336222.2) major capsid protein (*mcp*) gene by utilising the Stellaris® RNA FISH Probe Designer (Biosearch Technologies, Inc., Petaluma, CA) available online at www.biosearchtech.com/stellarisdesigner (Version 4.2; masking level=2, oligo length=20, minimum spacing=2). To avoid potential off-target binding we performed blastn (Altschul et al., 1990) searches of the 68 possible probe sequences against the *Acanthamoeba terricola Neff* host genome (RefSeq: GCF_000313135.1) and other giant virus genomes (*Tupanvirus deep ocean*: MF405918.1, *Tupanvirus soda lake*: KY523104.1, *Marseillevirus sp*. strain Vienna: PP736097.1, *Acanthamoeba castellanii medusavirus* J1: AP018495.1). Out of the 68 sequences, we selected 35 probes for smFISH (Table S1). The probe sequences were hybridised with the Quasar 670. Stellaris RNA FISH Probe set labeled with (Biosearch Technologies, Inc.), following the manufacturer’s instructions online at www.biosearchtech.com/stellarisprotocols.

### Sample preparation

*A. terricola Neff* (ATCC 30010) cells were seeded in a 24-well plate (Thermo Fischer Scientific #142475) in Peptone-Yeast Extract-Glucose medium (PYG: ATCC Medium 712) at a concentration of 10^5^ cells/mL and left to attach at 25°C for 30 min. *Acanthamoeba polyphaga mimivirus* was added at an MOI of 1, and the plates were centrifuged at 1000×g at room temperature (RT) for 30 min to synchronise the infection. To minimise the possibility of re-infections, the medium was exchanged after centrifugation.

After the desired hours post-infection (h.p.i), the cells were detached by scraping and the cell suspension of each well (∼1 mL) was collected in a 1.5 mL Eppendorf tube. The tubes were centrifuged at 5000×g at RT for 5 min, 800 μL of the supernatant was removed, and the pellet was resuspended in the remaining 200 μL through vortexing. Then, 50 μL of each cell suspension was added to a well of a microscope slide (Marienfeld #1216690) and the cells were left to attach for 1 hour at RT. The supernatant was removed from each well, and the samples were fixed with 20 μL 4% formaldehyde for 10 min at RT. Subsequently, the formaldehyde was removed, and the slides were washed with 40 μL of RNAse-free water. The cells were permeabilised with 40 μL of 70% ethanol for 1 hour at 4°C. After permeabilisation, 70% ethanol was removed from each well and further left to evaporate. To test whether the probes bind to RNA and not to DNA, an RNAse treatment was performed by adding a final concentration of 10 μg/mL RNAse A (Thermo Scientific #EN0531) for 30 min at 37°C to the slides before permeabilisation.

### Fluorescent noncanonical amino acid tagging (FUNCAT)

For performing FUNCAT, the amino acid homopropargylglycine (HPG) was added to the amoeba culture at least 30 min before sampling at a concentration of 50 µM. The dye mix (1.25 μL of 20 mM CuSO_4_, 1.25 μL of 100 mM THPTA and 0.3 µL 5 mM Cy3-azide) was prepared and incubated in the dark at RT for 3 min. Then, the dye mix was gently mixed with a freshly prepared mix of 221 μL Page’s Amoeba Saline buffer (PAS: ATCC medium 1323), 12.5 μL of 100 mM sodium ascorbate, and 12.5 μL of 100 mM aminoguanidine hydrochloride. Of this final mix, 10 μL was added to each well of the microscope slide. The slides were incubated in the dark at RT for 30 min. Afterwards, each well was washed 1-3 times with PAS.

### Single-molecule messenger RNA fluorescence in situ hybridisation (smFISH)

After permeabilisation, and optionally the FUNCAT click reaction, the slide was air-dried and 10 µL of smFISH Hybridisation Buffer (Vincent et al., 2021; 20% deionised formamide, 1 mg/mL *E. coli* tRNA, 2X SSC, 0.2 mg/mL BSA, 2 mM ribonucleoside-vanadyl complex, 100 mg/mL dextran sulfate, nuclease-free water) containing 437.5 nM of probe working solution was added to each well of the slide. For hybridisation, the slide was incubated in a moist chamber in the dark at 37°C overnight (∼20 hours). The smFISH Hybridisation Buffer was removed and the slide was transferred into a 50 mL tube containing Wash Buffer (2X SSC, Nuclease-free water) and incubated in a water bath at 39°C for 10 min. After washing, the slides were dipped into ice-cold Milli-Q water and air-dried. The different conditions tested for setting up the smFISH protocol and obtaining an optimal signal are displayed in Table 2.

**Table 2.** Conditions tested for performing the smFISH protocol.

|  |  |  |  |  |
| --- | --- | --- | --- | --- |
| Sample preparation | Fixation | 4% formaldehyde: 1 hour, 30 min, 10 min at RT |  |  |
|  | Permeabilisation | -70% Ethanol: 1 hour and 30 min at 4°C<br>-proteinase K: between 1 to 10 min at RT |  |  |
| Hybridisation |  | **smFISH** | **rRNA FISH** | **Stellaris RNA FISH** |
|  | Ingredients (final concentration) | 1.25, 2.5, 3.75, 4.375, 5 ng/μL probe mix |  |  |
|  |  | 0%, 10%, 15%, 20%, 25%, 30%, 35%, 40%, 50% formamide |  |  |
|  |  | 1 mg/mL *E. coli* tRNA | 900 mM NaCl | Stellaris Hybridization Buffer |
|  |  | 2X SSC (saline-sodium citrate) | 20 mM Tris/HCl |  |
|  |  | 0.2 mg/mL BSA (bovine serum albumin) | 0.01 % SDS |  |
|  |  | 2 mM ribonucleoside-v anadyl complex |  |  |
|  |  | 100 mg/mL dextran sulfate |  |  |
|  |  | Nuclease free water |  |  |
|  | Temperature | 32°C, 37°C, 42°C |  |  |
|  | Time | 5 hours, 17 hours, 20 hours |  |  |
| Post-hyb ridisation | Initial wash | 20% formamide, 2X SSC, 15 min at 37 °C and 39°C |  | Wash Buffer A |
|  | Final wash | Ice-cold Milli-Q water |  | Wash Buffer B |

### Ribosomal RNA fluorescence in situ hybridisation (rRNA FISH)

For rRNA FISH, 10 µL of rRNA FISH Hybridization Buffer (900 mM NaCl, 20 mM Tris-HCl pH 8.0, 0.01% SDS, and 25% formamide) containing 5 µM of Euk516 probe (5′-GGAGGGCAAGTCTGGT-3′; labelled with the fluorophore Fluos) working solution, was added to each well of the microscopy slide. Hybridisation was performed in a moist chamber in the dark at 46°C for 1.5 hours. Subsequently, slides were washed with a wash buffer (201 mM Tris-HCl pH 8.0, 50.25 mM EDTA, and 0.149 M NaCl) and kept in the wash buffer in a 50 mL tube at 48°C for 10 min. Then, the slides were dipped in ice-cold Milli-Q water and air-dried.

### DiD staining

DiD (1,1′-dioctadecyl-3,3,3′,3′-tetramethylindodicarbocyanine, 4-chlorobenzenesulfonate salt; Thermo Fischer Scientific #D7757) staining was performed after permeabilisation. The staining solution (1 µM DiD in 96-100% Ethanol) was sonicated for 30 min. After this homogenisation step, 3 µL of staining solution was added to each well of the microscope slide and incubated at 37°C for 20 min. After incubation, the wells were washed with 40 µL of PAS at RT for 5 min.

### Imaging

After the smFISH, FUNCAT, rRNA FISH and/or DiD staining, 4,6-diamidin-2-phenylindole (DAPI) (1 µg/mL) was added to the wells on the slides, incubated for 3 min, washed off with nuclease-free water, and air-dried. The slides were embedded in CitiFluor AF1 Mounting Medium (Electron Microscopy Sciences #17970-25) and imaged using a confocal laser-scanning microscope (Leica SP8).

### MOI experiments and digital polymerase chain reaction (dPCR)

*A. terricola Neff* cells were infected with *A. polyphaga mimivirus* (as described above) using an MOI of 0.5, 2, 5 and 10. Each well from a 24-well plate was sampled at 4 and 14 h.p.i, and for each MOI and time point, at least five different replicates (wells) were used, and the experiment was repeated 3 times, giving a minimum of 15 data points per MOI and time point.

For DNA extraction, the content of each well was collected in a 1.5 mL Eppendorf tube and centrifuged for 30 min at 13000×g at 10°C. The supernatant was replaced with 200 µL PAS, and the pellet was resuspended through vortexing. DNA was extracted using the DNeasy Blood & Tissue Kit (Qiagen #69506), adding 20 µL proteinase K and 200 µL Buffer AL and incubating at 56°C for 30 min at 400 rpm at the first steps. The DNA copy number of Mimivirus was quantified by performing digital PCR (dPCR) on the extracted DNA using the QIAcuity system and QIAcuity EG PCR Kit (Qiagen #250113). Primers targeting the *mcp* gene of Mimivirus (MCP_APolyphagaMimivirus_F: 5’-TCGTTTTTACGAAACATGATGG-3’; MCP_ApolyphagaMimivirus_R: 5’-CGATGGTGATTTGGAACACA-3’) were used as a proxy for measuring viral copy number.

For each MOI, the samples were also subjected to smFISH, DAPI staining and confocal laser-scanning microscopy. To count the number of eVFs at 4 h.p.i, at least 50 infected cells were used per MOI. To count the number of distinct *mcp* mRNA spots at 14 h.p.i, at least 45 infected cells were used per MOI.

### Statistics and data visualisation

The data was processed and visualised using R v.4.3.0 (R Core Team, 2023), with the packages dplyr (Wickham et al., 2023) and ggplot2 (Wickham, 2016). Final figures and graphs were made with Inkscape v.1.1.2 (https://inkscape.org).

## Data availability

The raw microscopy images and data for generating the figures can be found at Zenodo https://doi.org/10.5281/zenodo.15079575.

## Acknowledgements

We would like to thank Flora Vincent for the initial discussions on how to set up the smFISH protocol for *Nucleocytoviricota*. This project has received funding from the European Union’s Horizon 2020 research and innovation programme under the Marie Sklodowska-Curie grant agreement No. 891572 and the European Union (ERC, CHIMERA, 101039843). Views and opinions expressed are, however, those of the author(s) only and do not necessarily reflect those of the European Union or the European Research Council Executive Agency. Neither the European Union nor the granting authority can be held responsible for them. This research was funded in part by the Austrian Science Fund (FWF) [10.55776/COE7].

## References

1. Aagaard, K. M., Lahon, A., Suter, M. A., Arya, R. P., Seferovic, M. D., Vogt, M. B., Hu, M., Stossi, F., Mancini, M. A., Harris, R. A., Kahr, M., Eppes, C., Rac, M., Belfort, M. A., Park, C. S., Lacorazza, D., & Rico-Hesse, R. (2017). Primary Human Placental Trophoblasts are Permissive for Zika Virus (ZIKV) Replication. Scientific Reports, 7(1), 41389. 10.1038/srep41389

2. Altschul, S. F., Gish, W., Miller, W., Myers, E. W., & Lipman, D. J. (1990). Basic local alignment search tool. Journal of Molecular Biology, 215(3), 403–410. 10.1016/S0022-2836(05)80360-2

3. Aylward, F. O., Moniruzzaman, M., Ha, A. D., & Koonin, E. V. (2021). A phylogenomic framework for charting the diversity and evolution of giant viruses. PLOS Biology, 19(10), e3001430. 10.1371/journal.pbio.3001430

4. Dauty, E., & Verkman, A. S. (2005). Actin Cytoskeleton as the Principal Determinant of Size-dependent DNA Mobility in Cytoplasm: A NEW BARRIER FOR NON-VIRAL GENE DELIVERY*. Journal of Biological Chemistry, 280(9), 7823–7828. 10.1074/jbc.M412374200

5. Dieterich, D. C., Hodas, J. J. L., Gouzer, G., Shadrin, I. Y., Ngo, J. T., Triller, A., Tirrell, D. A., & Schuman, E. M. (2010). In situ visualization and dynamics of newly synthesized proteins in rat hippocampal neurons. Nature Neuroscience, 13(7), 897–905. 10.1038/nn.2580

6. Dieterich, D. C., Link, A. J., Graumann, J., Tirrell, D. A., & Schuman, E. M. (2006). Selective identification of newly synthesized proteins in mammalian cells using bioorthogonal noncanonical amino acid tagging (BONCAT). Proceedings of the National Academy of Sciences, 103(25), 9482–9487. 10.1073/pnas.0601637103

7. Hatzenpichler, R., & Orphan, V. J. (2016). Detection of Protein-Synthesizing Microorganisms in the Environment via Bioorthogonal Noncanonical Amino Acid Tagging (BONCAT). In T. J. McGenity, K. N. Timmis, & B. Nogales (Eds.), Hydrocarbon and Lipid Microbiology Protocols: Single-Cell and Single-Molecule Methods (pp. 145–157). Springer. 10.1007/8623_2015_61

8. Iwasaki, S., & Ingolia, N. T. (2017). The Growing Toolbox for Protein Synthesis Studies. Trends in Biochemical Sciences, 42(8), 612–624. 10.1016/j.tibs.2017.05.004

9. Koonin, E., Dolja, V., Krupovic, M., Varsani, A., Wolf, Y., Yutin, N., Zerbini, F., & Kuhn, J. (2019, October 14). Create a megataxonomic framework, filling all principal taxonomic ranks, for DNA viruses encoding vertical jelly roll-type major capsid proteins. ICTV Proposal: 2019.003G. 10.13140/RG.2.2.14886.47684

10. La Scola, B., Audic, S., Robert, C., Jungang, L., de Lamballerie, X., Drancourt, M., Birtles, R., Claverie, J.-M., & Raoult, D. (2003). A giant virus in amoebae. Science (New York, N.Y.), 299(5615), 2033. 10.1126/science.1081867

11. Legendre, M., Audic, S., Poirot, O., Hingamp, P., Seltzer, V., Byrne, D., Lartigue, A., Lescot, M., Bernadac, A., Poulain, J., Abergel, C., & Claverie, J.-M. (2010). mRNA deep sequencing reveals 75 new genes and a complex transcriptional landscape in Mimivirus. Genome Research, 20(5), 664–674. 10.1101/gr.102582.109

12. Minnigan, H., & Moyer, R. W. (1985). Intracellular location of rabbit poxvirus nucleic acid within infected cells as determined by in situ hybridization. Journal of Virology, 55(3), 634–643. 10.1128/jvi.55.3.634-643.1985

13. Mutsafi, Y., Shimoni, E., Shimon, A., & Minsky, A. (2013). Membrane Assembly during the Infection Cycle of the Giant Mimivirus. PLoS Pathogens, 9(5), e1003367. 10.1371/journal.ppat.1003367

14. Mutsafi, Y., Zauberman, N., Sabanay, I., & Minsky, A. (2010). Vaccinia-like cytoplasmic replication of the giant Mimivirus. Proceedings of the National Academy of Sciences, 107(13), 5978–5982. 10.1073/pnas.0912737107

15. Pasulka, A. L., Thamatrakoln, K., Kopf, S. H., Guan, Y., Poulos, B., Moradian, A., Sweredoski, M. J., Hess, S., Sullivan, M. B., Bidle, K. D., & Orphan, V. J. (2018). Interrogating marine virus-host interactions and elemental transfer with BONCAT and nanoSIMS-based methods. Environmental Microbiology, 20(2), 671–692. 10.1111/1462-2920.13996

16. Raj, A., van den Bogaard, P., Rifkin, S. A., van Oudenaarden, A., & Tyagi, S. (2008). Imaging individual mRNA molecules using multiple singly labeled probes. Nature Methods, 5(10), 877–879. 10.1038/nmeth.1253

17. Raoult, D., Audic, S., Robert, C., Abergel, C., Renesto, P., Ogata, H., La Scola, B., Suzan, M., & Claverie, J.-M. (2004). The 1.2-Megabase Genome Sequence of Mimivirus. Science, 306(5700), 1344–1350. 10.1126/science.1101485

18. Rigou, S., Schmitt, A., Lartigue, A., Danner, L., Giry, C., Trabelsi, F., Belmudes, L., Olivero-Deibe, N., Couté, Y., Berois, M., Legendre, M., Jeudy, S., Abergel, C., & Bisio, H. (2024). Nucleocytoviricota viral factories are transient organelles made by phase separation (p. 2024.09.01.610734). bioRxiv. 10.1101/2024.09.01.610734

19. Schramm, B., & Locker, J. K. (2005). Cytoplasmic Organization of POXvirus DNA Replication. Traffic, 6(10), 839–846. 10.1111/j.1600-0854.2005.00324.x

20. Vincent, F., Sheyn, U., Porat, Z., Schatz, D., & Vardi, A. (2021). Visualizing active viral infection reveals diverse cell fates in synchronized algal bloom demise. Proceedings of the National Academy of Sciences of the United States of America, 118(11), e2021586118. 10.1073/pnas.2021586118

21. Willemsen, A., Manzano-Marín, A., & Horn, M. (2024). Novel high-quality amoeba genomes reveal widespread codon usage mismatch between giant viruses and their hosts (p. 2024.09.23.614596). bioRxiv. 10.1101/2024.09.23.614596

22. Zauberman, N., Mutsafi, Y., Halevy, D. B., Shimoni, E., Klein, E., Xiao, C., Sun, S., & Minsky, A. (2008). Distinct DNA Exit and Packaging Portals in the Virus Acanthamoeba polyphaga mimivirus. PLoS Biology, 6(5), e114. 10.1371/journal.pbio.0060114

23. Zhang, R., Mayer, L., Hikida, H., Shichino, Y., Mito, M., Willemsen, A., Iwasaki, S., & Ogata, H. (2024). Giant virus creates subcellular environment to overcome codon– tRNA mismatch (p. 2024.10.07.616867). bioRxiv. 10.1101/2024.10.07.616867

