## Supplementary material for "Giant virus transcription and translation occur at well-defined locations within amoeba host cells": Fig. S1-S5

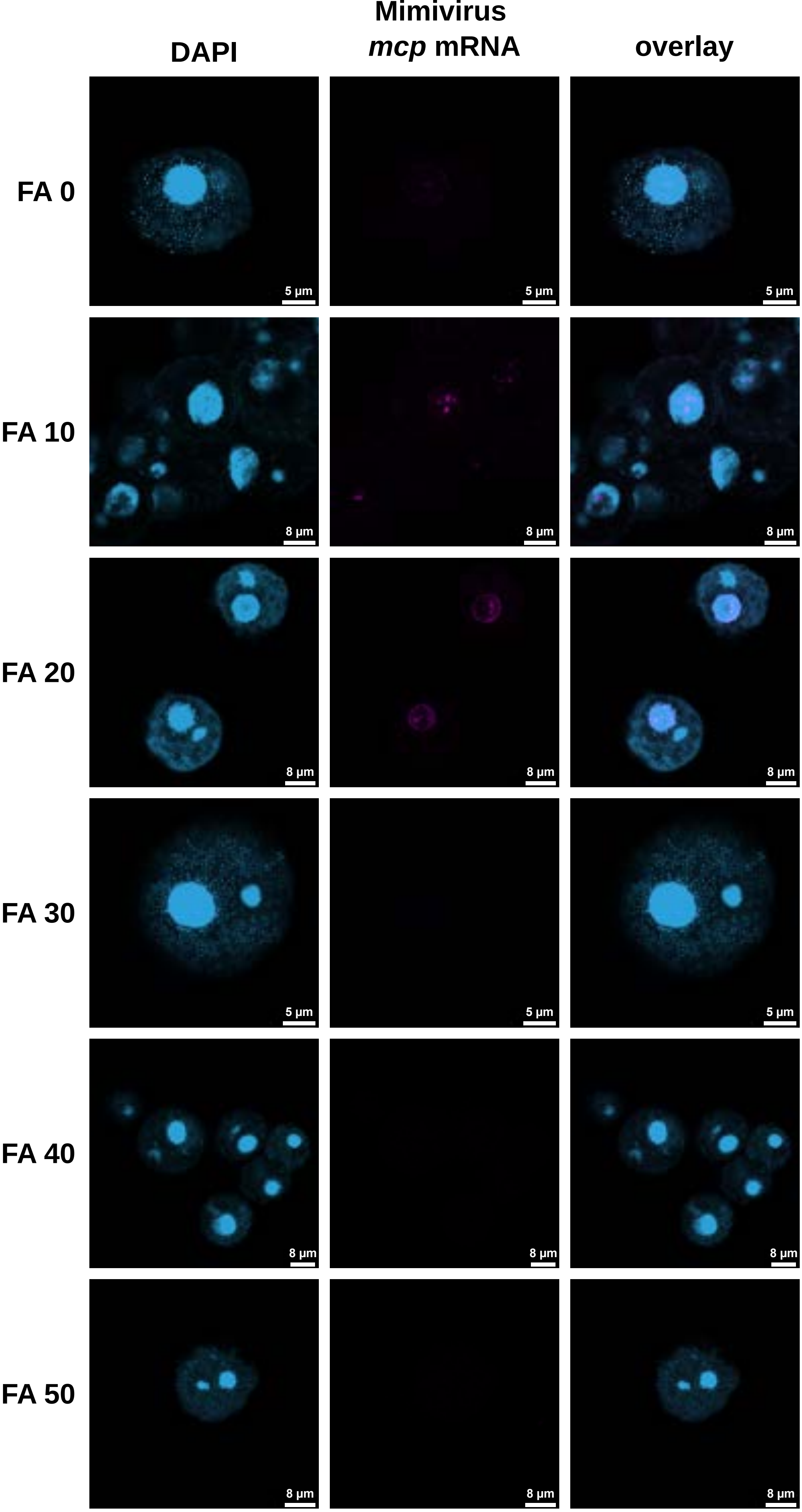


**Figure S1**. **Formamide series**. Different formamide (FA) concentrations (0-50%) in the hybridisation buffer were used to determine the optimal concentration for the probe mix and target sequence. The first column is the signal of DAPI staining (light blue), the middle column is the Mimivirus *mcp* mRNA signal (magenta), and the third column is the overlay.

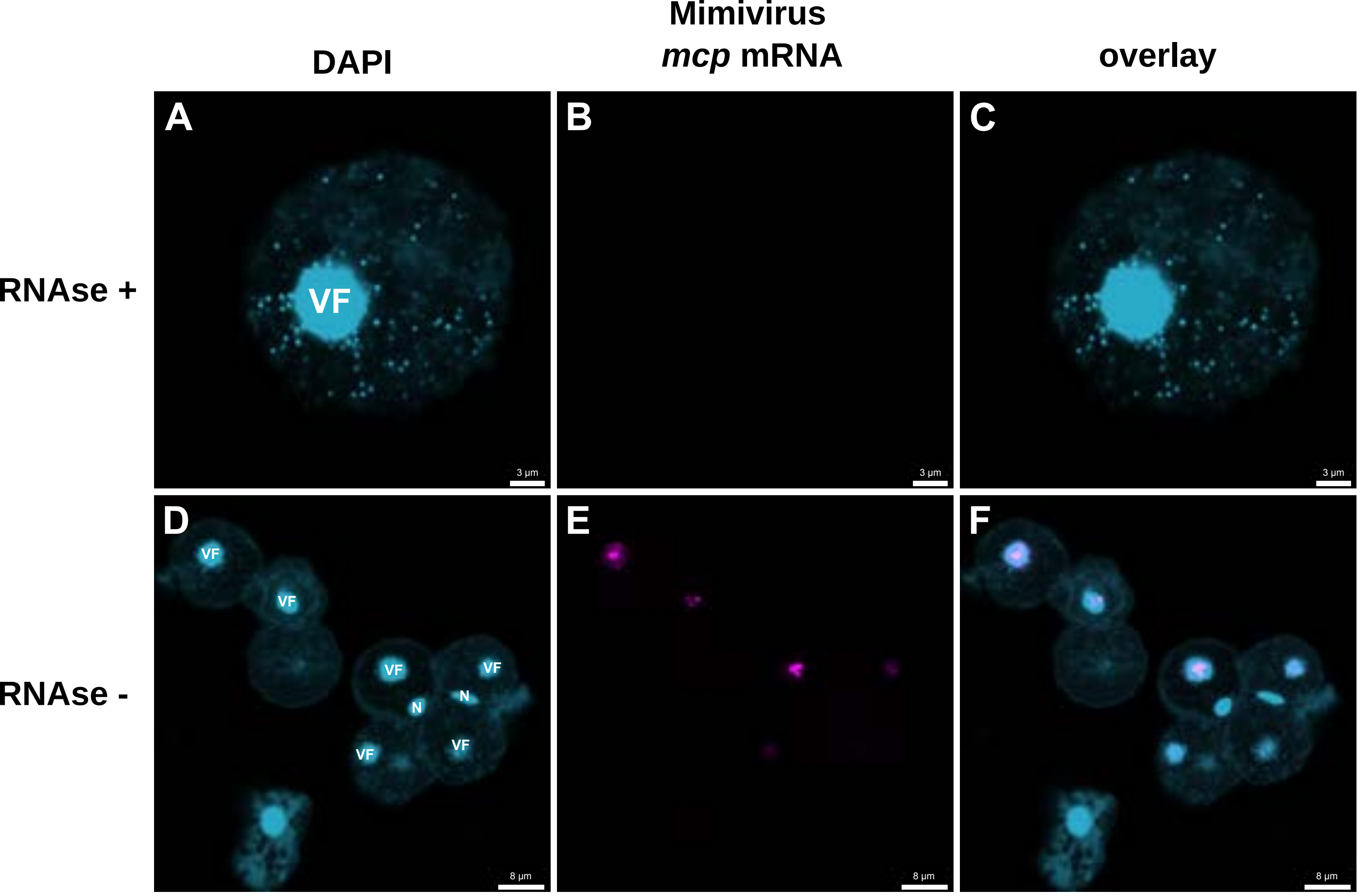


**Figure S2**. **RNAse control**. To confirm that the probes bind to RNA and not DNA, the cells were treated with RNAse A (**A**, **B**, **C**) and compared to the untreated condition (**D**, **E**, **F**). DAPI staining (light blue), Mimivirus *mcp* mRNA signal (magenta), and the overlay of infected amoeba cells.

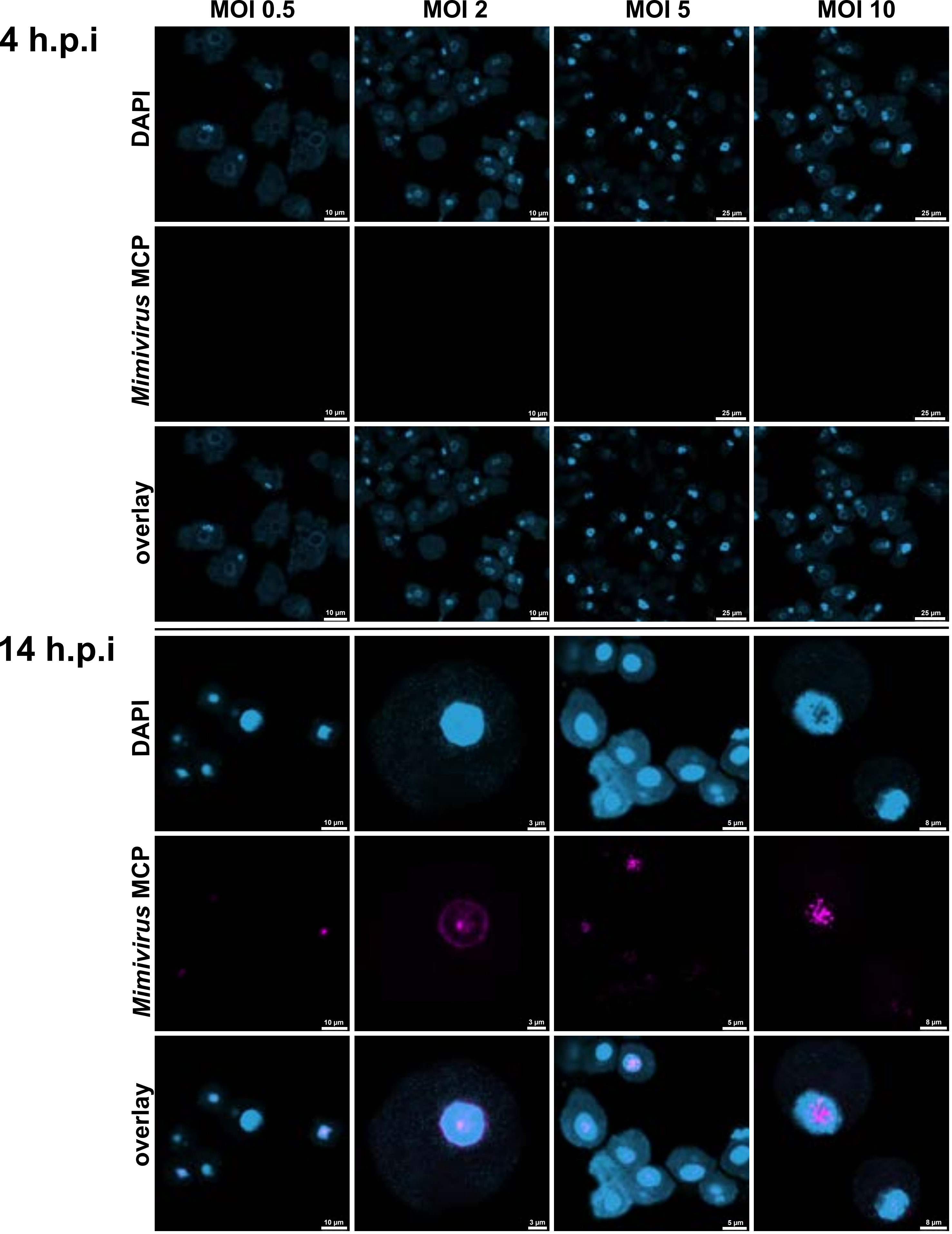


**Figure S3**. **The full version of Fig. 3A**. DAPI staining (light blue) and Mimivirus *mcp* mRNA signal (magenta) of amoeba cells infected with different MOIs at 4 and 14 h.p.i. The Mimivirus *mcp* mRNA signal can not be observed at 4 h.p.i as expression levels are undetectable at this time point.

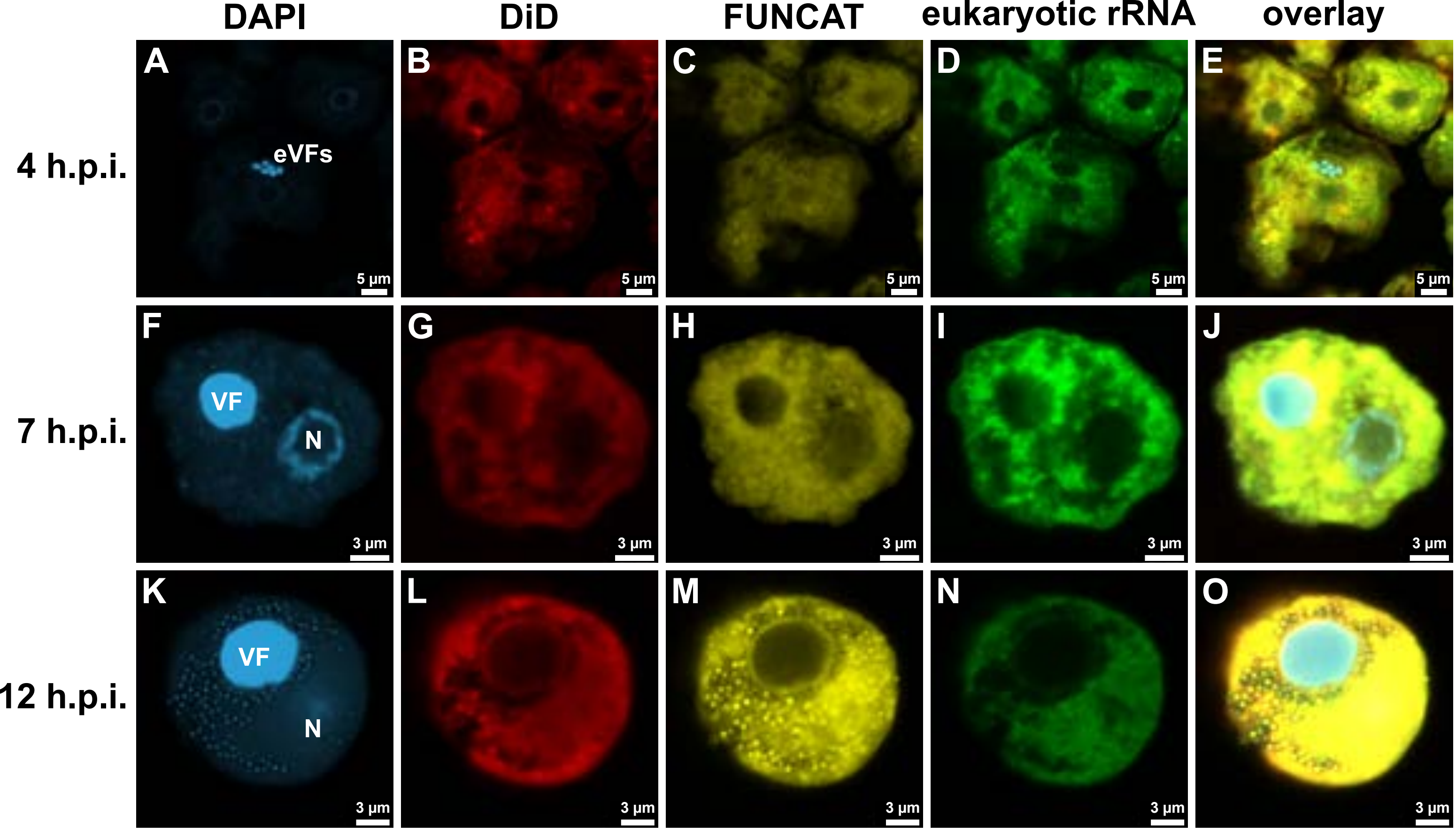


**Figure S4. Different labelling techniques support the idea that Mimivirus translation occurs in a ring surrounding the viral factory**. DAPI staining (**A**, **F**, **K**), DiD labelling (**B**, **G**, **L**), FUNCAT (**C**, **H**, **M**), rRNA FISH (**D**, **I**, **N**), and an overlay (**E**, **J**, **O**) of amoeba host cells infected with Mimivirus at 4 (**A**-**E**), 7 (**F**-**J**), and 12 (**K**-**O**) h.p.i. DAPI staining (light blue) reveals multiple early viral factories (eVFs) at an early infection stage (**A**, **E**). At later stages of infection, eVFs fuse to become mature VFs (**F**, **J**, **K**, **O**). A low DiD signal (red) can be observed both at the nucleus and at the site of eVFs and VFs (**B**, **G**, **L**), while a higher signal can be observed in a ring surrounding the VF (**L**). FUNCAT (yellow) also reveals a strong signal in a ring surrounding the VF (**M**). The rRNA FISH signal (green) also accumulates in a ring surrounding the VF (**N**).

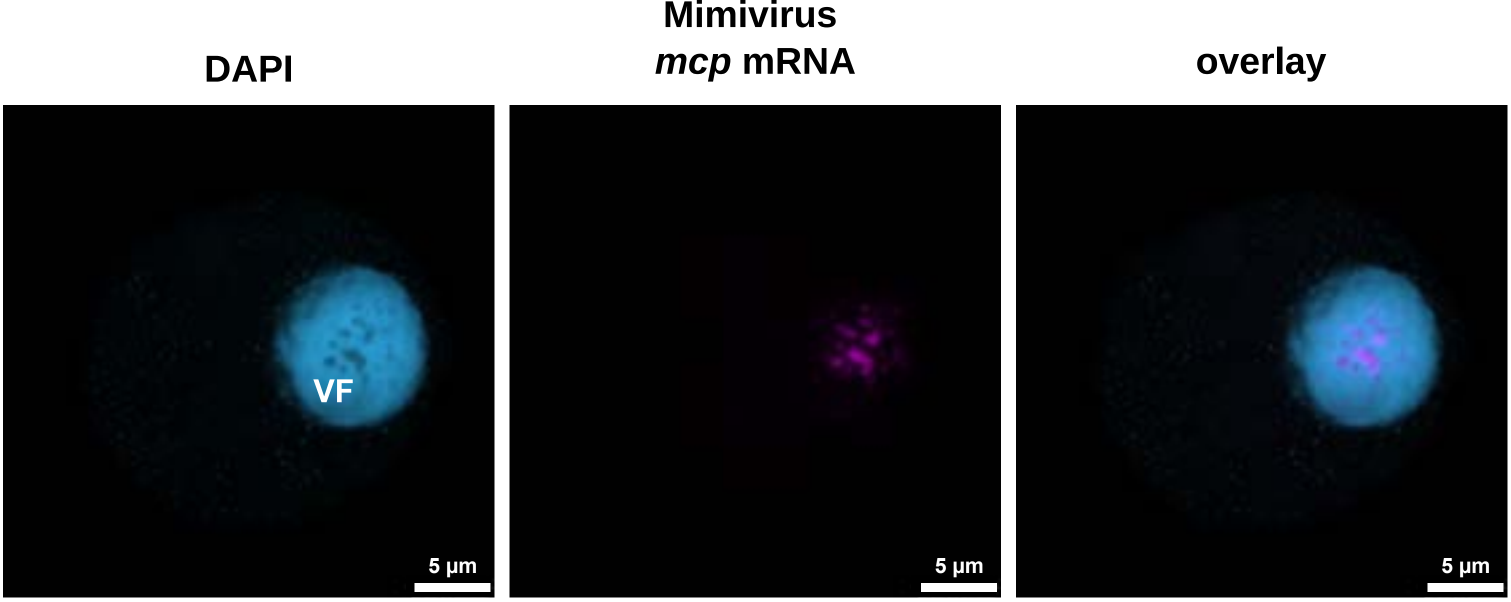


**Figure S5**. **Image supporting the idea of specific transcription sites**. DAPI staining (light blue) and Mimivirus *mcp* mRNA signal (magenta) of one infected amoeba cell. This image shows a lack of DAPI signal within the VF at the spots where Mimivirus *mcp* mRNA is present.
